# Topical Ophthalmic Administration of VIAN-c4551 Antiangiogenic Peptide for Diabetic Macular Edema: Preclinical Efficacy and Ocular Pharmacokinetics

**DOI:** 10.1101/2024.09.11.612517

**Authors:** Elva Adán-Castro, Magdalena Zamora, Daniela Granados-Carrasco, Lourdes Siqueiros-Márquez, Jose F. García-Rodrigo, Thomas Bertsch, Jakob Triebel, Gonzalo Martínez de la Escalera, Juan Pablo Robles, Carmen Clapp

## Abstract

VIAN-c4551 is a cyclic antiangiogenic peptide that stands as a potent and stable inhibitor of vascular endothelial cell growth factor (VEGF), the major vasopermeability and angiogenic factor in diabetic macular edema and diabetic retinopathy. Intravitreal injections of inhibitors of VEGF are a first-line therapy, but the invasiveness, risk, and low adherence of frequent intravitreal injections interfere with the needed long-term treatments and successful outcomes. Eye drops are non-invasive and favor compliance. Here, we evaluated the preclinical efficacy, permeability, and ocular pharmacokinetics of VIAN-c4551 delivered in eye drops. VIAN-c4551 demonstrated high potency (IC_50_ = 137 pM) to inhibit the permeability of human umbilical vein endothelial cell monolayers induced by VEGF. VIAN-c4551 eye drops potently (0.005% minimum effective dose) prevented the retinal vascular leakage induced by VEGF injected intravitreally for up to 24 hours and reversed the increase in retinal vascular permeability due to streptozotocin-induced diabetes in rats and mice. VIAN-c4551 exhibited high permeability across MDCK epithelium and, after a single topical ocular instillation in rabbits, reached the retina-choroid in micromolar concentrations several orders of magnitude above its IC_50_ (C_max_= 51 µM at 6 hours) that lasted at least 24 hours. In conclusion, VIAN-c4551 eye drops reach the back of the eye at therapeutic concentrations, providing a potential, once-a-day, non-invasive intervention for preventing and reversing retinal vascular leakage in diabetic macular edema, diabetic retinopathy, and other vascular retinopathies and preserving sight.

## Introduction

Diabetes is a common cause of vision loss due to retinal microvascular complications in diabetic retinopathy (DR) and diabetic macular edema (DME). DME is present in 25% of diabetics and 71% of patients with proliferative DR, and it is the clinical condition most closely associated with vision loss.^1^ In DME, leakage from retinal capillaries causes the accumulation of extracellular fluid and proteins that alter the structure and function of the macula and can lead to blindness if untreated. Vascular endothelial growth factor (VEGF) is a major vascular permeability factor in DME,^2^ and the intravitreal injection of agents blocking VEGF has become the first-line treatment for DME with vision loss.^3^ Nevertheless, intravitreal injections are invasive, and their repeated and long-term use, due to the chronic nature of DME, increases the chance of ocular complications and systemic adverse events.^4^ Eye drops are a preferred therapeutic option as these can be self-administered, are non-invasive, and facilitate long-term therapy. However, reaching the back of the eye through eye drops is challenging due to the numerous ocular barriers,^5,6^ and no eye-drop formulation has been approved by the FDA to treat posterior eye diseases, including DME.

VIAN-c4551 is a promising drug candidate for treating DME, DR, and other vascular retinopathies. VIAN-c4551 is an analog of vasoinhibin, an endogenous antiangiogenic protein with significant therapeutic potential. Vasoinhibin inhibits endothelial cell proliferation, migration, survival, and permeability in response to VEGF and other proangiogenic and vascular permeability factors (basic fibroblast growth factor, bradykinin, interleukin 1β).^7,8^ It helps restrict retinal angiogenesis under physiological conditions,^9^ and the intravitreal administration of vasoinhibin reduces ischemia-induced retinal angiogenesis^10^ and prevents and reverses diabetes- and VEGF-induced increase in retinal vascular leakage in rodents.^11^ In humans, the systemic and ocular levels of vasoinhibin are disrupted in retinopathy of prematurity^12,13^ and DR,^14^ and a prokinetic medication (levosulpiride) that upregulates vasoinhibin levels in the vitreous of patients with proliferative DR,^15^ improves visual acuity and structural retinal outcomes in patients with DME.^16^

However, the clinical use of vasoinhibin is hampered by difficulties in its recombinant production.^17^ These difficulties have been overcome by the development of VIAN-c4551, a vasoinhibin analog with improved pharmacological properties that conserves the efficacy and potency of vasoinhibin.^8^ VIAN-c4551 inhibits VEGF-induced proliferation of endothelial cells with potency like vasoinhibin (IC_50_=150 pM) and is orally active to inhibit melanoma tumor growth and vascularization in mice.^8^ The oral bioavailability of VIAN-c4551 demonstrates physiochemical properties (solubility, stability, permeability, and potency) that may facilitate VIAN-c4551 to overcome the anatomical and physiological barriers that prevent topically administered drugs from reaching the retina.^5,6^ Here, we show the high potency and efficacy of VIAN-c4551 in eye drops to inhibit VEGF- and diabetes-induced retinal vascular leakage in rodents, the in vitro permeability of VIAN-c4551 across epithelial cells, and the presence of VIAN-c4551 in the retina-choroid and vitreous at supra-therapeutic concentrations following eye drop administration in rats and rabbits.

## Material and methods

### Reagents

VIAN-c4551(>95% pure), VIAN-c4551 coupled to fluorescein-5-isothiocyanate (FITC), and recombinant human vascular endothelial growth factor-165 (VEGF) were from GenScript (Piscataway, NJ). Basic fibroblast growth factor (bFGF) from Scios, Inc., Mountain View, CA. Recombinant human vasoinhibin of 123 amino acids (Vi1-123) was produced as reported.^18^ Eye drop vehicle (3% trehalose−0.15% sodium hyaluronate) was from Laboratories Théa (Clermont-Ferrand, France).

### Cell culture

Human umbilical vein endothelial cells (HUVEC)^19^ were cultured in F12K medium supplemented with 20% fetal bovine serum (FBS), 100 µg mL^-1^ heparin (Sigma Aldrich), 25 µg mL^-1^ endothelial cell growth supplement (ECGS) (Corning, Glendale, AZ), and 100 U mL^-1^ penicillin-streptomycin, and authenticated by CD31 presence. The Madin-Darby canine kidney (MDCK) epithelial cell line (ATCC, Rockville, MD) was cultured in 10% FBS−DMEM.

### VEGF-induced endothelial cell permeability

HUVEC cells were seeded in 20% FBS−F12K−heparin-ECGS on a 6.5 mm transwell with a 0.4 µm pore to form a confluent monolayer. Transendothelial electrical resistance (TEER) was measured with the EVOM^2^ Volt/Ohm meter (World Precision Instruments, Sarasota, FL), as reported.^8^ Monolayers were incubated with different concentrations of VIAN-c4551 or Vi1-123 for 1 hour before adding PBS or 50 ng mL^-1^ VEGF, and TEER measured over a 150-min period.

### MDCK permeability assay

MDCK cells were seeded (40,000 cells cm^-2^) in 200 µL 10% FBS−DMEM on a 6.5 mm transwell with a 0.4 µm pore to form a confluent monolayer. Upon confluency, 0.4 nM and 4 nM VIAN-c4551 or Vi1-123 were added to the apical side of the monolayer for 2 hours at 37^°^C. 600 μL from the basolateral compartments were then collected and stored at -20^°^C. VIAN-c4551 and Vi1-123 were quantified by their capacity to inhibit HUVEC proliferation as previously reported.^20^ Recovery levels were relative to those without MDCK cells, and the apparent permeability coefficient (*P*_*app*_, cm s^-1^) was calculated as reported.^21^

### Animals

Male and female Wistar rats (6−7 weeks old, 180−230 g), male CD-1 mice (8−10 weeks old, 20−25 g), and female New Zealand white rabbits (2-2.5 kg) were maintained and treated in adherence to the National Research Council’s Guide for the Care and Use of Laboratory Animals. The Bioethics Committee of the Instituto de Neurobiología (UNAM) approved all animal experiments.

### VEGF-induced retinal vascular permeability

One drop containing 0.5% [5 mg mL^-1^ or 6.5 mM] VIAN-c4551 or vehicle was instilled in each eye of mice (5 µL) and rats (10 µL) manually maintained still for 10 seconds to allow drop distribution on the ocular surface. After 1 hour, animals were anesthetized with 60% ketamine and 40% xylazine (1 µL g^-1^ body wt, i.p.) and injected intravitreally with 1 μL (mice) or 2 µL (rats) PBS or VEGF (250 ng), and Evans blue dye (45 mg/kg) injected intravenously. Two hours later, retinal vascular leakage was evaluated by the extravasation of albumin as reported.^22^

### Diabetes-induced retinal vascular permeability

Five daily i.p. injections of streptozotocin (STZ) (55 mg/kg; S0130, Sigma–Aldrich) and a single i.p. STZ injection (60 mg/kg) induced diabetes in male mice and male rats, respectively.^23,24^ Blood glucose levels >250 mg dL^-1^ defined diabetes one week post-STZ. After 5 weeks, non-diabetic and diabetic animals received a daily eye drop of 0.5% VIAN-c4551 or vehicle per eye for 5 days. We performed the Evans blue method 24 hours after the last eye drop.

### Ocular pharmacokinetics

Female rats and rabbits were instilled with one eye drop (10 and 50 µL, respectively) containing 0.5% VIAN-c4551, VIAN-c4551-FITC (net peptide), or vehicle, and manually kept still for 10 seconds to allow drop surface distribution and euthanized by CO^2^ inhalation at different intervals for up to 24 hours. Eyes were enucleated, and the vitreous, the retina from the rat, and the retina-choroid from the rabbit were collected, snap frozen in liquid nitrogen, and stored at -80^°^C until analysis. Tissues were homogenized in RIPA buffer (10 mM Tris-HCl, 140 mM NaCl, 0.5% IGEPAL CA-630, 1mM EDTA, 0.5 mM EGTA, 1% triton X-100, 0.1% SDS, pH 7.4) supplemented with 1:100 protease inhibitor cocktail (Roche). Vitreous samples (125 ng μL^-1^ protein) and retinas (150 ng μL^-1^ protein) were added to HUVEC cultures to evaluate VIAN-c4551 levels through its bioactivity, as reported.^20^ For fluorescence emission, a standard curve of VIAN-c4551-FITC from 0 to 100 pmol was prepared in a vitreous matrix (110 ng μL^-1^ protein) for rat or rabbit or in a retina matrix (2.5 μg μL^-1^ protein for rat or 750 ng μL^-1^ protein for rabbit). Fluorescence emission of 100 µL from each standard dilution was quantified (Varioskan Flash, Thermo Fisher Scientific, Waltham, MA) at excitation and emission wavelengths of 490 nm and 520 nm, respectively, for 100 ms. Values were interpolated from standard curves in the respective matrix and pharmacokinetic analysis was performed using Plotly software.^25^

### Statistical analysis

Statistical analysis was performed using the GraphPad Prism version 10.2.2 for macOS, GraphPad Software (Boston, Massachusetts USA, www.graphpad.com). The threshold for significance was set at *P*<0.05.

## Results

### VIAN-c4551 potently inhibits the VEGF-induced permeability of HUVEC monolayers

We evaluated the trans-endothelial electrical resistance (TEER) of HUVEC monolayers as an indicator of vascular permeability (Figure 1). VEGF rapidly decreased TEER values and kept them low throughout the 150-minute incubation period. VIAN-c4551 prevented the VEGF-induced reduction of TEER, and VIAN-c4551 alone showed no effect (Figure 1A). TEER inhibition by VIAN-c4551 was dose-dependent with an IC_50_ of 137.3 pM, like that of the full-length vasoinhibin (Vi1-123) (IC_50_ = 163.2 pM) (Figure 1B).

**Figure 1.**
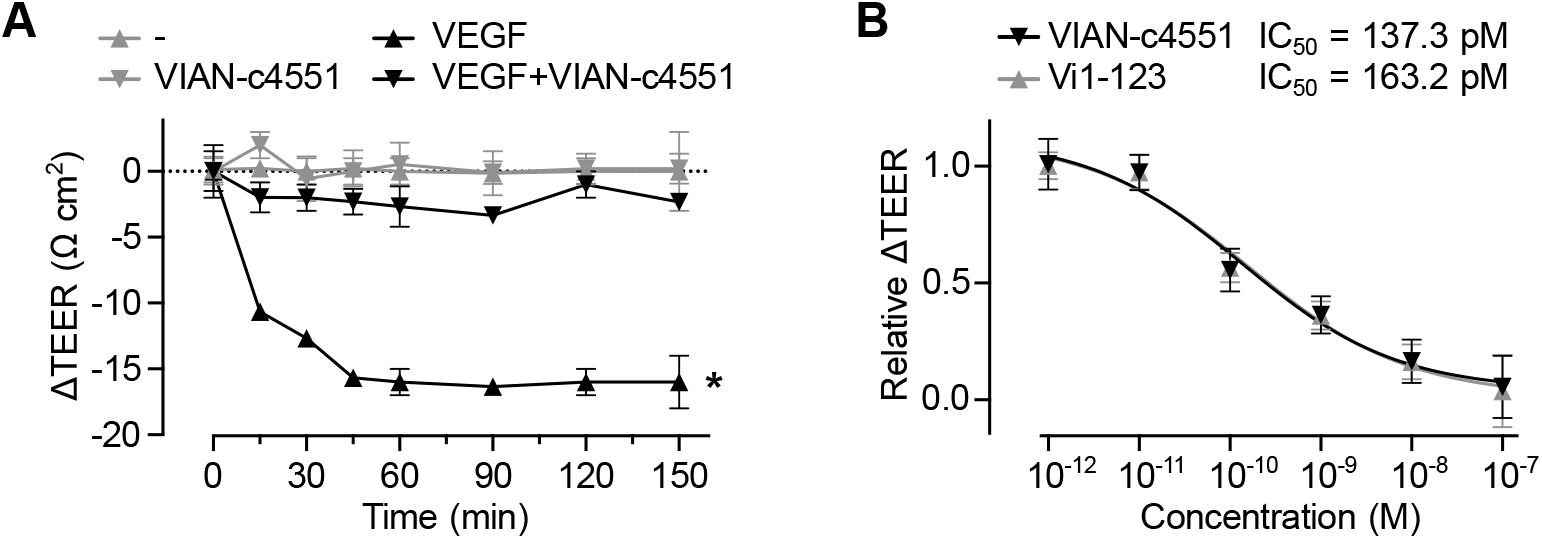
VIAN-c4551 inhibits VEGF-induced permeability of HUVEC monolayers. (A) Changes in the trans-endothelial electrical resistance (ΔTEER) of HUVEC monolayers treated or not with 100 nM VIAN-c4551 throughout a 150-minute incubation period without (-) or with VEGF (50 ng mL^-1^). Values are means ± SD, n=3, \**P*<0.006 vs. (-) for all time points except 0 (2-way ANOVA repeated measures, Dunnett’s). (B) Dose-response inhibition of the VEGF-induced reduction of ΔTEER at 90 minutes by different concentrations of VIAN-c4551 or Vi1-123. The IC_50_ of VIAN-c4551 and Vi1-123 are indicated. ΔTEER is relative to that of VEGF alone. Values are means ± SD, n=9, and the curve represents a nonlinear regression analysis of four parameters (54 points, r^2^ ≥ 0.92).

### VIAN-c4551 exhibits high permeability across epithelia

We evaluated the capacity of VIAN-c4551 to cross the MDCK epithelial cell monolayer. Around 50% of VIAN-c4551 in two concentrations (0.4 and 4 nM) crossed the monolayer after 2 hours (Figure 2A). VIAN-c4551 presented a high apparent permeability coefficient (*P*_*app*_ = 1.44 × 10^−5^ cm s^-1^) (Figure 2B), like steroids.^26^ The permeability of Vi1-123 across the MDCK monolayer was negligible (*P*_*app*_ = 3.86 × 10^−7^ cm s^-1^).

**Figure 2.**
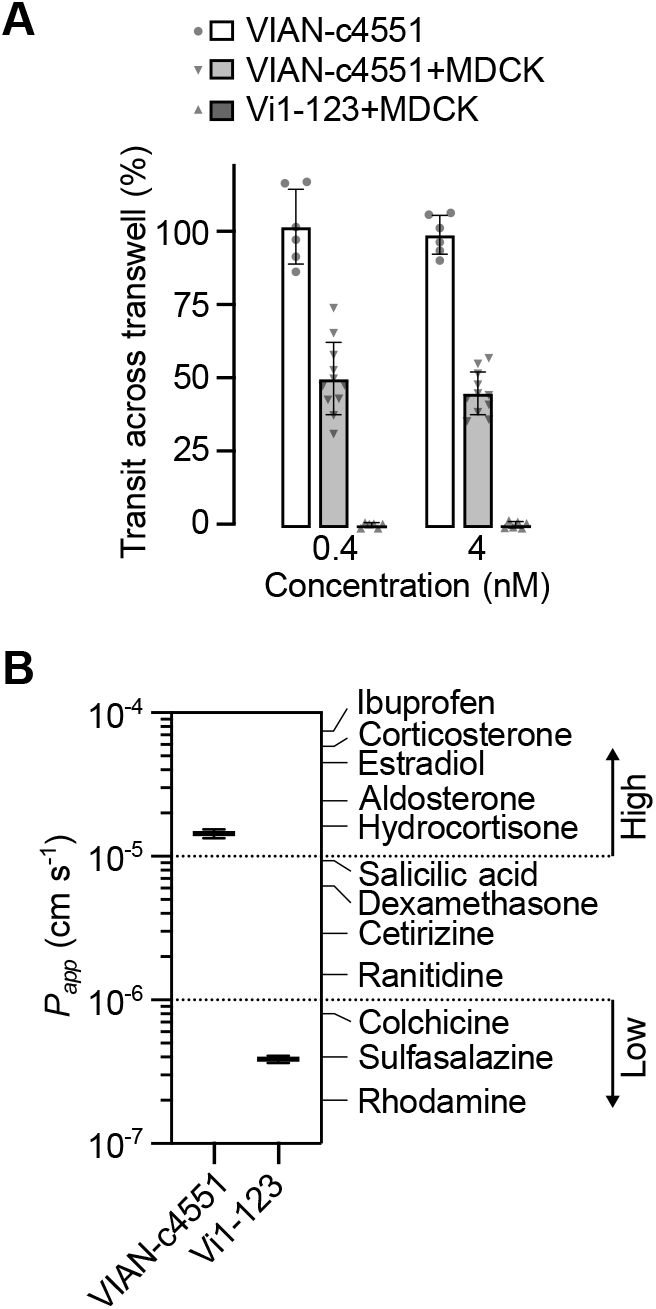
VIAN-c4551 permeability across MDCK epithelial cells. (A) Percentage of transit across an MDCK monolayer of two concentrations (0.4 and 4 nM) of VIAN-c4551 or Vi1-123 after 2 hours relative to their transit of both compounds in the absence of MDCK monolayers. Bars indicate mean ± SD, n ≥ 6. Individual values are shown. (B) Permeability coefficient (*P*_*app*_) of VIAN-c4551 and Vi1-123. The *P*_*app*_ of other drugs across MDCK monolayers are added as reference.^26^ Pointed lines indicate established limits between high and low permeability values established with reference drugs.

Given that VIAN-c4551 is highly potent to inhibit VEGF-induced vascular permeability and highly permeable across epithelia, we investigated the effect of VIAN-c4551 in eye drops on VEGF-induced retinal vascular permeability.

### VIAN-c4551 eye drops prevent VEGF-induced retinal vascular permeability

A single eye drop containing 0.5% VIAN-c4551 or vehicle was administered one hour before intravitreal injection of VEGF or PBS, and two hours later the retinal vascular leakage was evaluated by the extravasation of albumin. VEGF induced a significant 2–fold increase in retinal vascular permeability relative to PBS, and VIAN-c4551 prevented this effect in both rats (Figure 3A) and mice (Figure 3B). VIAN-c4551 did not modify the retinal levels of the tracer in the absence of VEGF.

**Figure 3.**
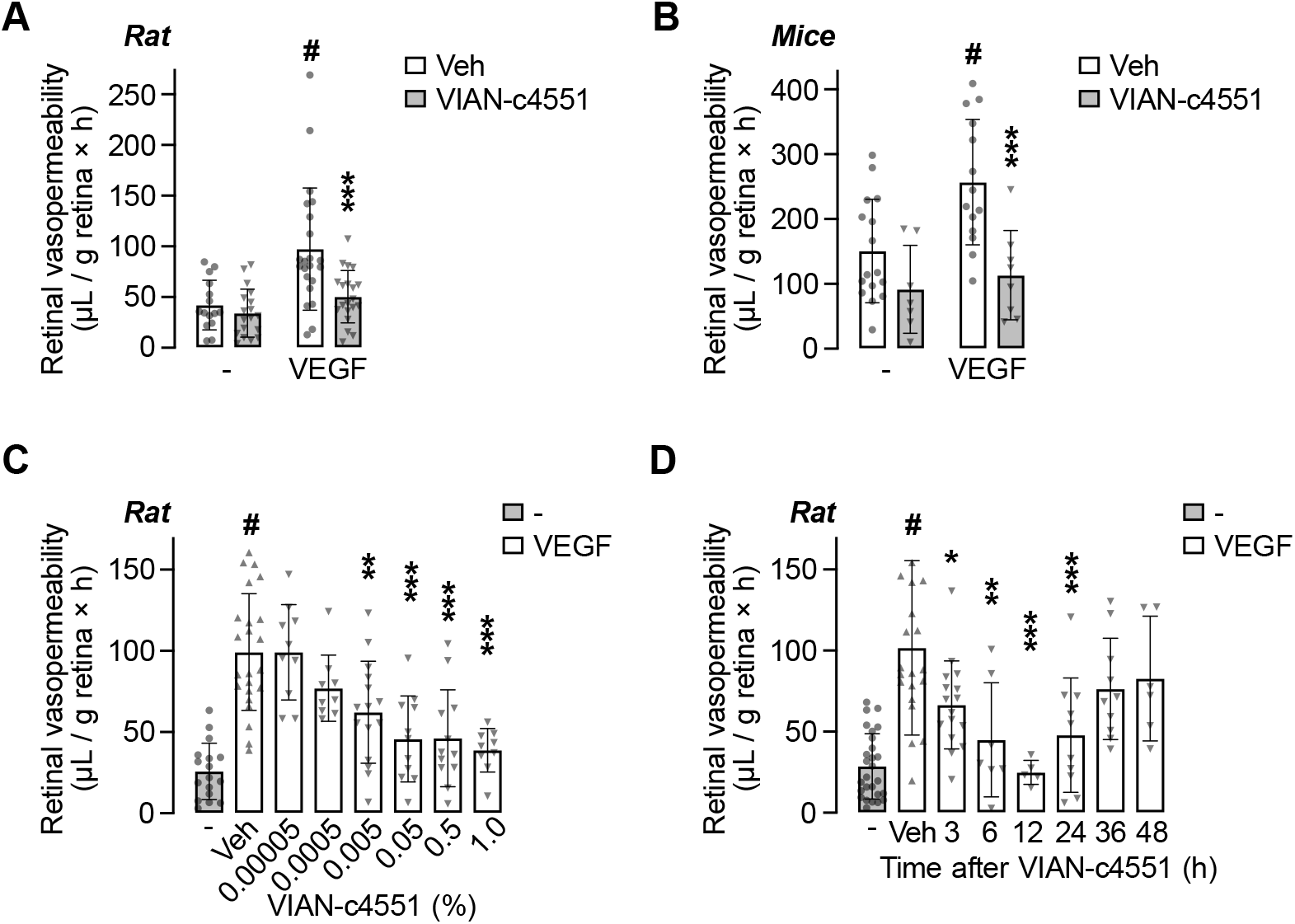
VIAN-c4551 eye drops potently inhibit VEGF-induced retinal vascular permeability for up to 24h. Retinal vascular permeability in rats (A) and mice (B) treated with a single eye drop containing 0.5% VIAN-c4551 or vehicle administered one hour before the intravitreal injection of VEGF or PBS. (C) Dose-response inhibition of eye drops containing different concentrations of VIAN-c4551. (D) Duration of the effect of a single 0.5% VIAN-c4551 eye drop administered at different times before the intravitreal injection of VEGF. Quantification of Evans blue-stained albumin extravasation was performed two hours after VEGF administration. Values are means ± SD. Individual eye values are shown. #*P*<0.002 vs no VEGF (-), \**P*<0.033, \*\**P*<0.002, \*\*\**P*<0.001 vs. Veh (-), 2-way ANOVA, Šidák’s (A, B), 1-way ANOVA, Dunnett’s (C, D).

The efficacy of VIAN-c4551, profiled in rats, was dose-dependent, and the minimal effective dose was 0.005% (0.5 µg/eye drop) (Figure 3C), confirming the high potency of VIAN-c4551. The duration of the effect was also profiled in rats by instilling 0.5% VIAN-c4551 or vehicle at different times before intravitreal injection of VEGF. The protective effect of VIAN-c4551 against the VEGF challenge was evident at 3 hours, maximal at 12 hours, and lasted up to 24 hours after the eye drop (Figure 3D). The exceptional efficacy and 24–hour duration of VIAN-c4551 in eye drops suggested that VIAN-c4551 reaches the back of the eye at a high and sustained therapeutic concentration and warranted investigating its effect in the DME and DR rodent model induced by STZ, and its pharmacokinetic profile.

### VIAN-c4551 in eye drops reverses diabetes-induced retinal vascular permeability

Excessive retinal vascular leakage is evident at 6 weeks of STZ-induced diabetes in rodents (Figure 4).^23^ A daily eye drop containing 0.5% VIAN-c4551 for the last five days suppressed the 2-fold increase in retinal vascular permeability due to diabetes in rats (Figure 4A) and mice (Figure 4B). The reduction of vascular leakage is indicative of the restoration of the blood-retinal barrier function and, thereby, proof-of-concept that VIAN-c4551 represents a potential, non-invasive therapy against DR and DME.

**Figure 4.**
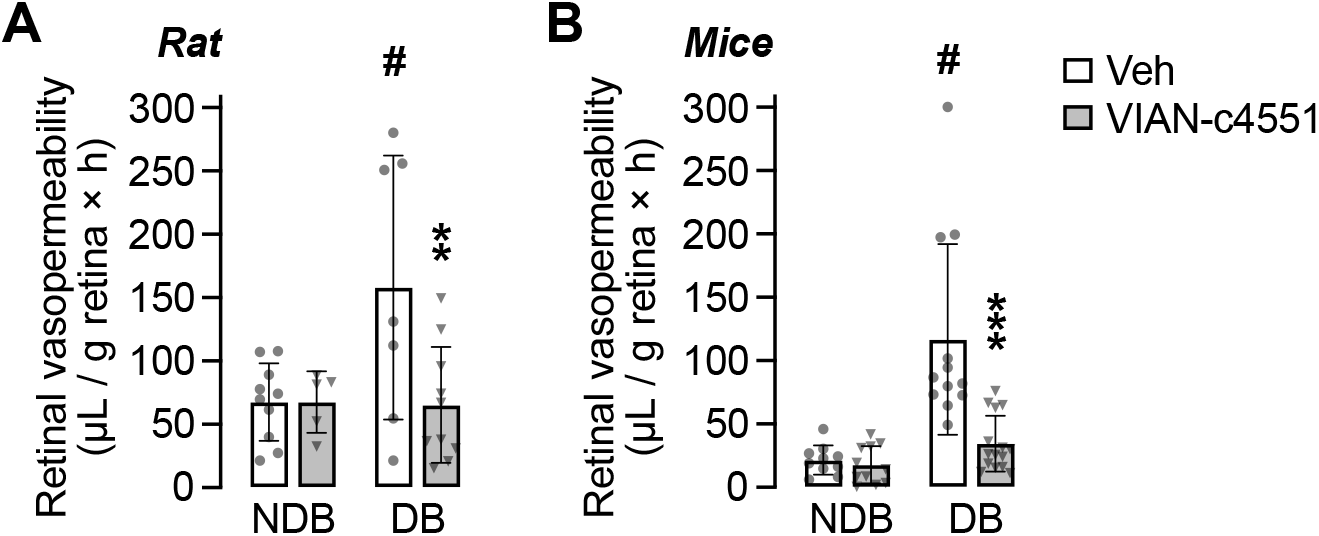
VIAN-c4551 inhibits diabetes-induced retinal vascular permeability. Retinal vascular permeability in nondiabetic (NDB) and 6-week diabetic (DB) rats (A) and mice (B) were treated daily for 5 days with a single eye drop containing 0.5% VIAN-c4551 or vehicle. The extravasation of Evans blue-stained albumin evaluated retinal vascular permeability 24 hours after the last eye drop. Values are means ± SD. Individual eye values are shown. #*P*<0.002 vs NDB, \*\**P*<0.002, \*\*\**P*<0.001 vs. Veh, (2-way ANOVA, Šidák’s).

### Topical administration of VIAN-c4551 leads to high and sustained therapeutic concentration at the back of the eye

After a single eye drop, VIAN-c4551 reached the retina at levels ∼300-fold greater than in the vitreous, and both amounts substantially exceeded the IC_50_ for inhibition of endothelial permeability. In rats, VIAN-c4551’s concentration in the retina was 32.8 µg g^-1^ (∼43 µM, assuming a tissue density of 1 g mL^-1^) and in the vitreous 300 ng mL^-1^ (∼391 nM) at 3 hours post-dose (Figure 5A). In rabbits, the pharmacokinetic (PK) profile defined a long absorption phase that peaked in the retina-choroid (C_max_ 52 µg g^-1^ or ∼67 μM) and in the vitreous (C_max_ 184 ng g^-1^ or ∼239 nM) at 6 hours and slightly declined thereafter (Figure 5B). Even after 24 hours, VIAN-c4551 exposure in retina-choroid (3.5 µg g^-1^ or ∼4.64 μM) and vitreous (10.6 ng mL^-1^or ∼13.88 nM) were >4 and >2 orders of magnitude higher than its IC_50_. PK parameters are summarized in Table 1. The PK profile was confirmed with VIAN-c4551-FITC. Despite that the FITC labeling reduced the permeability of VIAN-c4551 across MDCK monolayers (Figure 5C) and lowered its penetration to the retina-choroid and vitreous of rabbits (Figure 5D), the kinetic profile of VIAN-c4551-FITC in both compartments was like that of the unlabeled compound, i.e., C_max_ in retina-choroid was 1.3 µg g^-1^ (∼1.1 μM) at a T_max_ of 6 hours and maintained for 24 hours at levels beyond IC_50_.

**Table 1.** Ocular PK parameters after a single eyedrop of 0.5% VIAN-c4551 to NZ rabbits.

| PK Parameters | Retina-Choroid | Vitreous |
| --- | --- | --- |
| $C_{\text{max}}$ (ng g<sup>-1</sup> or ng mL<sup>-1</sup>) | 51,693.81 ± 6678.9 | 183.66 ± 15.49 |
| $T_{\text{max}}$ (hours) | 6 | 6 |
| AUC<sub>0-24h</sub> (hours × ng g<sup>-1</sup> or hours × ng mL<sup>-1</sup>) | 467,434 ± 45,884 | 1,678 ± 182 |
| $T_{1/2}$ (hours) | 5 ± 0.78 | 4.7 ± 0.58 |

Values are mean ± SEM. N = 3 eyes per timepoint. ng g<sup>-1</sup> is used for retina-choroid and ng mL<sup>-1</sup> for vitreous. AUC, area under the curve.

**Figure 5.**
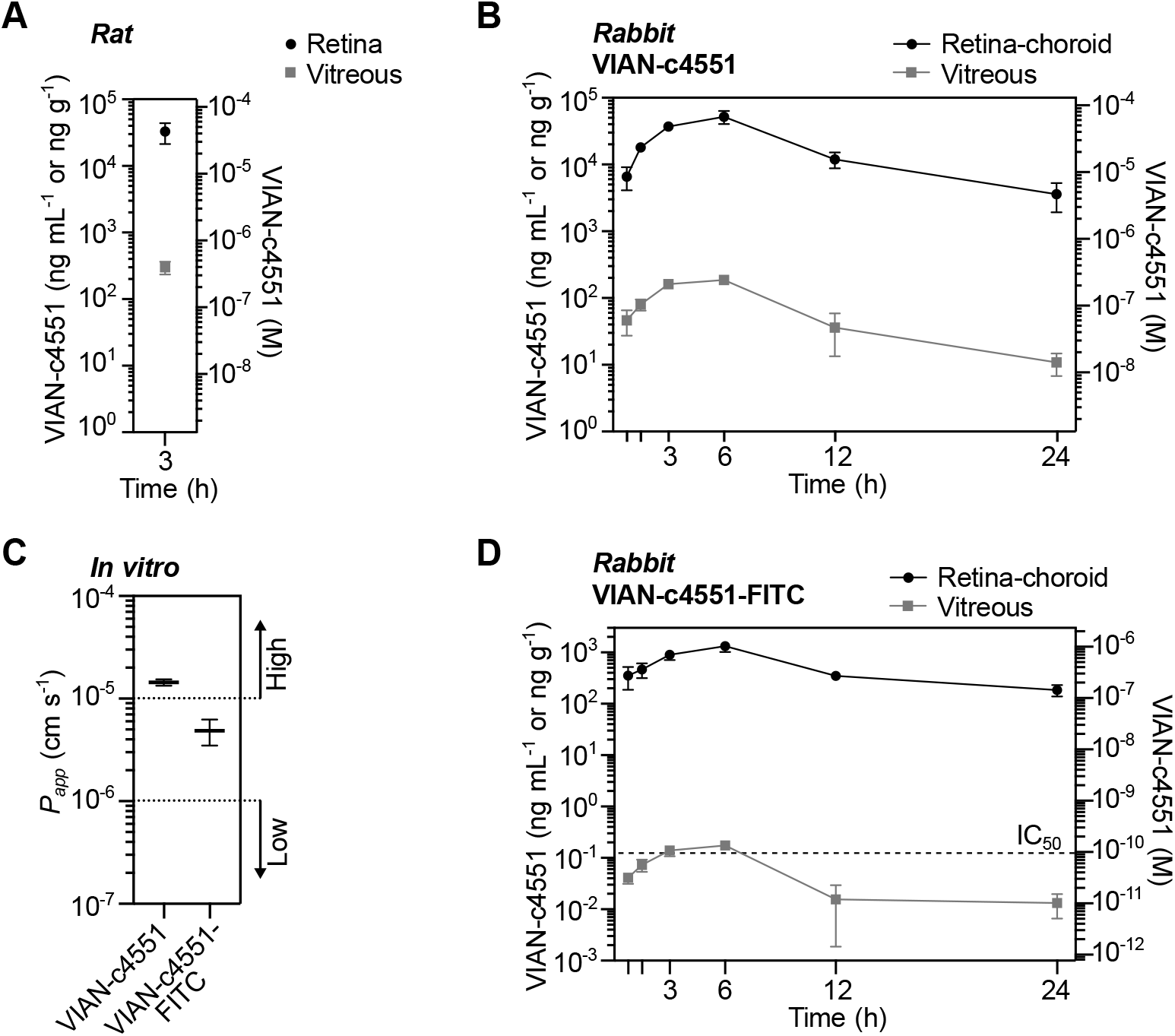
Ocular penetration and pharmacokinetics of VIAN-c4551. Concentration of VIAN-c4551 in the retina and vitreous of rats three hours after delivery of a single eye drop containing 0.5% VIAN-c4551 (A), and in the retina-choroid and vitreous of NZ rabbits at different time points after a 0.5% VIAN-c4551 eye drop (B). Values are mean ± SD, n ≥3; ng g^-1^ is used for retina-choroid and ng mL^-1^ for vitreous, and the right y-axis indicates molarity. (C) Permeability coefficient (*P*_*app*_) values of VIAN-c4551-FITC across MDCK epithelial cell monolayer compared to that of VIAN-c4551. Pointed lines indicate established limits between high and low permeability values established with reference drugs. (D) Concentration of VIAN-c4551-FITC in the retina-choroid and vitreous of rabbits at different time points after 0.5% VIAN-c4551-FITC eye drop instillation. Values are mean ± SD, n ≥3; ng g^-1^ is used for retina-choroid and ng mL^-1^ for vitreous, and the right y-axis indicates molarity.

## Discussion

Excessive vascular leakage is a hallmark of retinal diseases with high prevalence, such as DME, DR, and age-related macular degeneration. The current therapy uses frequent anti-VEGF intravitreal injections, which are highly invasive, not always effective, cause complications, and associate with low compliance. Eye drop delivery is much more appealing as it is applied non-invasively by the patients themselves. However, topical administration faces delivery problems since only very low concentrations of drugs reach the retina.^6,27,28^ Here, we report the eye drop delivery of VIAN-c4551, a small peptide derived from an endogenous protein, that has outstanding anti-VEGF efficacy and pharmacokinetic properties and has the potential to eliminate the need for repeated intravitreal injections.

VIAN-c4551 was developed as an effort to translate the anti-angiogenic vasoinhibin family of proteins (5.6 –18 kDa) into an alternative therapeutic compound of 768 Da with improved pharmacological properties. In VIAN-c4551, the functional domain (His46-Gly47-Arg48) of vasoinhibin is contained within its 7 amino acid sequence (residues 45 to 51 of vasoinhibin) synthesized with D–amino acids in reverse order and cyclized to conserve potency and improve stability. VIAN-c4551 has the same vasoinhibin potency (IC_50_ = 150 pM) to inhibit VEGF-induced endothelial cell proliferation and is orally active to inhibit melanoma tumor growth and vascularization in mice.^8^ Furthermore, the apoptotic, inflammatory, and fibrinolytic effects of vasoinhibin, which may be undesirable for antiangiogenic therapy, are in a different structural domain (residues 30 to 36) and, thereby, absent in VIAN-c4551.^29^

Here, we report for the first time the inhibitory effect of VIAN-c4551 on excessive vascular permeability, another vascular action of vasoinhibin.^11,24^ VIAN-c4551 blocked the VEGF-induced permeability of endothelial cell monolayers with the high potency (IC_50_=137 pM) of vasoinhibin. Such high potency is key for topical delivery to inhibit excessive retinal vascular leakage, as VIAN-c4551 needs to cross several ocular barriers and reach the retina at therapeutic concentrations. Overcoming ocular barriers depends on physicochemical properties, including size, charge, lipophilicity, solubility in water, stability, and permeability.^5,27^ The oral bioavailability of VIAN-c4551 is indicative of its capacity to cross the intestinal system into the bloodstream and, thereby, of having adequate stability and permeability across epithelial barriers.

Aiming to develop the eye drop delivery of VIAN-c4551, we investigated its permeability value across MDCK cell epithelial monolayers. The MDCK cell system has been proposed as a model for predicting the penetration of drugs across the cornea^30^ and for studying the outer blood-retinal barrier.^31^ VIAN-c4551 presented a high apparent permeability coefficient in MDCK cells, with values of magnitude over 10^−5^ cm s^-1^ and an average recovery of more than 50%. Therefore, VIAN-c4551 is highly permeable, and its transport across ocular epithelial barriers (corneal, conjunctiva, sclera, choroid, and retinal pigment epithelium) may not represent a relevant restriction for its topical absorption.

Encouraged by its high anti-VEGF potency and high permeability, our efforts turned toward testing the in vivo efficacy of VIAN-c4551 in eye drops against the intravitreal injection of VEGF, a non-diabetic model of DME and DR^32^. A single eye drop containing 0.5% VIAN-c4551 prevented the increase in retinal vascular permeability induced by VEGF in rats and mice. The effect was dose-dependent and highly potent as the minimal effective dose was 0.005% (0.5 µg/eye/day). This dose is much lower than used for the topical delivery of antibodies (7.5 mg/eye/day), antibody fragments (0.2 − 0.5 mg/eye/day), or small molecules (0.3% − 1% /eye/day) in preclinical models of retinal microvascular diseases or clinical trials.^27,33,34^ Moreover, the anti-VEGF effect of VIAN-c4551 lasted 24 hours, suggesting a successful once-a-day topical ocular treatment.

VEGF is a major contributor to retinal vascular leakage in diabetes.^2^ However, the pathologic cascade leading to DME and DR also involves other vasoactive substances like bradykinin,^35^ bFGF,^36^ and interleukin-1β,^37^ all of which are inhibited by vasoinhibin.^7^ By interacting with binding proteins (plasminogen activator inhibitor, urokinase, the urokinase receptor, and/or integrin α5β1)^38,39^ in endothelial cell membranes, vasoinhibin inhibits signaling pathways (Ras-Raf-MAPK, Ras-Tiam1-Rac1-Pac1, PI3K-Akt, and PLCγ-IP3-eNOS) activated by several proangiogenic and vascular permeability factors (VEGF, bFGF, bradykinin, and interleukin β1).^7^ Accordingly, vasoinhibin inhibits the increase in retinal vascular permeability induced by the intravitreal injection of vitreous from patients with DR,^40^ and the ocular overexpression of vasoinhibin is more efficient than that of sFlt-1 (the secreted extracellular domain of the VEGF receptor 1) for blocking diabetes-induced retinal vascular permeability.^24^ Addressing only the inhibition of VEGF action may, therefore, be of limited value, and the efficacy of VIAN-c4551 in diabetic conditions could exceed that of agents targeting only VEGF.

To investigate the broader benefit of the topical application of VIAN-c4551, we used the pharmacological induction of diabetes with STZ in rodents, one of the most used DME and DR experimental models. In the STZ model of diabetes, increased retinal vascular permeability can occur as early as after 2 weeks of hyperglycemia.^32^ Daily eye drops of 0.5% VIAN-c4551 for 5 days suppressed the retinal vascular leakage caused by 6 weeks of hyperglycemia in rats and mice. This result indicated that VIAN-c4551 in eye drops reverses the disruption of the blood-retinal barrier in diabetic rodents and, thereby, has potential curative properties in DME, DR, and other microvascular retinal diseases. The efficacy of VIAN-c4551 in the VEGF-non-diabetic model and the STZ-diabetic model is proof-of-concept that its topical delivery reaches the retina at therapeutic concentrations.

In agreement, VIAN-c4551 efficiently migrates to the back of the eye of rats and rabbits. A single 0.5% VIAN-c4551 eyedrop reached micromolar and nanomolar concentrations in the retina/choroid and vitreous, respectively, and the high levels were sustained for 24 hours. The maximal concentrations in the retina/choroid (C_max_ ∼67 µM) and vitreous (C_max_ ∼239 nM) were 5 and 3 orders of magnitude higher, respectively, than the VIAN-c4551 IC_50_ (137.3 pM) to inhibit VEGF-induced endothelial cell permeability. These levels position VIAN-c4551 as the instilled compound with the highest measured concentration in the retina of rabbits (reviewed here^6^), the next one being dorzolamide hydrochloride, a glaucoma drug yielding a 2.8 µM retinal concentration after a 2% instillation dose.^6^ However, pooling together the choroid and retina limits comparisons as it is difficult to conclude exposure in the retina. Also, albino animals confound the more general condition of pigmented animals, where binding to melanin can interfere with the ocular efficacy and distribution of drugs.^41^

A major question relates to the mechanisms mediating the exceptional ocular pharmacokinetic properties of VIAN-c4551. The fact that the concentration of VIAN-c4551 is lower in the vitreous than in the retina/choroid speaks in favor of migration through the conjunctiva-sclera-choroid-retina route and from the retina to the vitreous. This possibility is consistent with the hydrophilic nature of VIAN-c4551 and with absorption of hydrophilic compounds being facilitated by the larger paracellular pores and surface area of the non-corneal route.^27^ However, VIAN-c4551 could also be absorbed through the cornea or through its clearance by the conjunctival blood vessels and subsequent retinal access via the retinal vasculature. Studies evaluating the ocular and systemic distribution of VIAN-c4551 following topical and intravenous delivery should help clarify these mechanisms. The physicochemical properties of VIAN-c4551 should also be addressed. Its small size, solubility in water, high permeability, and resistance to proteolysis very likely contribute to its efficient intraocular migration and 5-hour half-life in the inner ocular tissues.

In summary, our findings suggest the feasibility of the topical self-administration of VIAN-c4551 in an outpatient setting that would represent a breakthrough in ophthalmology. Currently, no topical ocular formulations are available for treating microvascular retinal diseases. Several small molecules have undergone clinical trials but have not proceeded to the market despite having successful preclinical data.^6,42^ The many reasons for failure include assuming the direct translation of positive results in rodents and rabbits to patients, which is limited by the anatomical and physiological differences between their eyes and the human eye.^27,43,44^ However, the high concentrations of VIAN-c4551 achieved in the posterior segment of the eye indicate that its topical use for treating retinal diseases may be possible. This possibility is further acceptable due to the nature of VIAN-c4551 as an analog of vasoinhibin, an endogenous protein restraining the vascularity of ocular tissues^9,45^ that is downregulated in retinopathy of prematurity,^12^ DR,^14^ and DME.^15,16^ Detailed physicochemical characterization, evaluation of safety profiles, and incorporation of large-scale production requirements warrant further research.

## Acknowledgments

The authors thank Fernando López Barrera, Xarubet Ruíz Herrera, Adriana González Gallardo, Alejandra Castilla, María A. Carbajo, and Martín García Servín for their excellent technical assistance.

## Notes

**Funding:** This work was supported by the Secretaría de Educación, Ciencia, Tecnología e Innovación de la Ciudad de México (SECTEI/061/2023).

### Competing Interest Statement

Funding: This work was supported by the Secretaria de Educacion, Ciencia, Tecnologia e Innovacion de la Ciudad de Mexico (SECTEI/061/2023).
Declaration of Competing Interests: JPR, MZ, TB, JT, GME, and CC are inventors of a submitted patent application (WO/2021/098996), which is owned by the Universidad Nacional Autonoma de Mexico (UNAM) and JT and TB. JPR is the CEO and founder of VIAN Therapeutics Inc. MZ and CC are consultants for VIAN Therapeutics. Inc.

